# Glioma CpG Island Methylator Phenotype (G-CIMP): Biological and Clinical Implications

**DOI:** 10.1101/169680

**Authors:** Tathiane M Malta, Camila F de Souza, Thais S Sabedot, Tiago C Silva, Maritza QS Mosella, Steven N Kalkanis, James Snyder, Ana Valeria B Castro, Houtan Noushmehr

## Abstract

Gliomas are a heterogeneous group of brain tumors with distinct biological and clinical properties. Despite advances in surgical techniques and clinical regimens, treatment of high-grade glioma remains challenging and carries dismal rates of therapeutic success and overall survival. Challenges include the molecular complexity of gliomas, as well as inconsistencies in histopathological grading, resulting in an inaccurate prediction of disease progression and failure in the use of standard therapy. The updated 2016 World Health Organization (WHO) classification of tumors of the central nervous system reflects a refinement of tumor diagnostics by integrating the genotypic and phenotypic features, thereby narrowing the defined subgroups. The new classification recommends the molecular diagnosis of the *IDH* mutational status in gliomas. *IDH*-mutant gliomas manifest the CpG Island Methylator Phenotype (G-CIMP). Notably, the recent identification of clinically relevant subsets of G-CIMP tumors (G-CIMP-high and G-CIMP-low) provide a further refinement in glioma classification that is independent of grade and histology. This scheme may be useful for predicting patient outcome and may be translated into effective therapeutic strategies tailored to each patient. In this review, we highlight the evolution of our understanding of the G-CIMP subsets and how recent advances in characterizing the genome and epigenome of gliomas may influence future basic and translational research.

**Conflict of interest:** The authors declare no conflict of interest

**ABBREVIATIONS**
2-HG: 2-hydroxyglutarate (2-HG)
α-KG: α-ketoglutarate
CNS: Central nervous system
CGI: CpG Island
CpG: 5'-C-phosphate-G-3'
CTCF: CCCTC-binding factor
G-CIMP: Glioma CpG island methylator phenotype
GBM: Glioblastoma (World Health Organization grade IV)
GWAS: Genome-wide association study
HDAC: Histone deacetylase
IDH: Isocitrate dehydrogenase
LGG: Lower-grade glioma (here defined by diffusely infiltrative low-grade or intermediate-grade glioma (World Health Organization grade II or III))
MGMT: O-6-methylguanine DNA methyltransferase
NAD: Nicotinamide adenine dinucleotide
OS: Overall survival
TMZ: Temozolomide
WHO: World Health Organization

## Gliomas: An Overview

Gliomas are a heterogenous group of brain tumors with distinct biological and clinical properties ^1,2^. Classically, glioma subtypes and grading were exclusively defined by histological features. However, this classification strategy does not reflect the heterogeneity of the tumor, is prone to subjectivity and discordance among neuropathologists, falls short of predicting disease course and cannot reliably guide treatment^2^. Therefore, multiple research efforts have sought to identify molecular signatures that define more discrete glioma subgroups and have a greater impact and relevance in the clinical setting ^1-5^. Corroborating the importance of molecular markers in the cancer classification, researchers observed that 1 in 10 cancer patients would be better classified by molecular taxonomy, rather than by the current system based on the primary tissue of origin and stage of disease ^6^.

The 2016 update to the WHO classification of tumors of the central nervous system (CNS) represents a shift in tumor diagnostics by integrating molecular and phenotypic features into the classification of tumors and thereby narrowing the defined subgroups ^1^. Among the genetic alterations associated with diffuse gliomas, *IDH* mutation, histone H3 K27M status, and the integrity status of chromosomes 1 and 19 (1p and 19q) are now integrated with the traditional histology/grade-based glioma classification ^1,7^ Gliomas harboring mutations in *IDH1/2*, as well as codeletion of the 1p and 19q chromosome arms (1p/19q), have shown favorable prognostic and/or predictive values in relation to their counterparts (IDH-wildtype or 1p/19q intact [also known as “non-codel”]) ^8-12^. A number of epigenomic markers also have shown prognostic and/or predictive values ^13^. Patients harboring gliomas that carry *MGMT* promoter DNA methylation demonstrate increased overall survival (OS) and time to progression of the disease after treatment with temozolomide (TMZ) or radiation ^14-17^. *MGMT* promoter DNA methylation status is used to guide therapeutic management for anaplastic gliomas with wildtype *IDH1/2* and in glioblastoma (GBM) of the elderly ^14-17^. Another important milestone highlighting the clinical importance of epigenetic signatures in gliomas was the discovery of the Glioma-CpG Island Methylator Phenotype (G-CIMP) ^18^. Patients carrying G-CIMP (G-CIMP positive) tumors have shown a better prognosis than those not carrying that phenotype (G-CIMP negative). G-CIMP positive (G-CIMP+) tumors were closely related to *IDH*-mutation and nearly all *IDH*-mutant gliomas were G-CIMP+ and had a favorable prognosis ^18,19^. However, comprehensive DNA methylation profiling in a large cohort of glioma patients revealed that not all *IDH*-mutant/G-CIMP+ tumors had the same prognosis ^7^. Based on the extent of global DNA methylation, this study uncovered two subsets of *IDH*-mutant/G-CIMP+ gliomas, one that presented a low degree of DNA methylation and poorer outcome (G-CIMP-low) and another subset depicting high DNA methylation and good OS as usually described for *IDH*-mutant/G-CIMP+ (G-CIMP-high) ^7,18,19^ G-CIMP+ subsets (-low and -high) have shown distinct biological features and clinical implications that will be further explored in this review. Following a brief introduction to the role of epigenomics in cancer and in neuro-oncology, we will detail the evolution of our understanding that led to the identification of the G-CIMP subsets. We will also describe how recent advances in DNA methylation-based biomarkers that characterize gliomas may influence future basic and translational research. Our review will conclude with current and upcoming clinical applications, focusing on the potential translational importance of G-CIMP subsets.

### Epigenetic Modifications in Cancer

Within the nucleus of a cell, the three billion base pairs of the human DNA sequence are stored and organized as chromatin, which is made up of repeating units called nucleosomes. Nucleosomes are formed by octamers of histone proteins that are prone to chemical modifications at distinct amino acid residues, such as methylation and acetylation. Several epigenetic mechanisms may operate in synchrony to modify the packaging of the genome, including DNA methylation, histone modifications, nucleosome remodeling, small and long non-coding RNA, protein:DNA interactions via chromatin modifying transcription factors, and three-dimensional chromatin architecture ^20^. The resulting epigenomic landscape determines the accessibility of regulatory elements, thereby modulating the transcriptional regulation given by transcription factors ^20^. Epigenetic mechanisms are implicated both in physiological and pathological events, such as tissue specificity and carcinogenesis, respectively ^20^. In the process of malignant transformation, gene inactivating mutations have been shown to control the epigenomic landscape, implying a crosstalk between the genome and the epigenome ^21^. In the present review, we will focus on the first epigenetic modification to be linked to cancer and probably the most extensively studied epigenetic modification in mammals - DNA methylation ^22^.

### DNA methylation

DNA methylation status results from the action of methylating or demethylating enzymes. DNA methylation is the covalent transfer of methyl groups to the 5' position of the cytosine ring, primarily at dinucleotide cytosine-p-guanines (CpGs), resulting in 5-methylcytosine (5mC). DNA methyltransferases (DNMTs), known as the DNA methylation “writer” enzymes, catalyze the transfer of the methyl group to 5’ cytosine ^23^. On the other hand, demethylation of 5mC is promoted by the TET1/2 (Ten-Eleven Translocation) methylcytosine dioxygenases, known as the DNA methylation “erasers”, to generate 5-hydroxymethylcytosine (5hmC) ^24^. Some stretches of DNA contain frequent CpG sites, defined as CpG islands (CGI), which are preferentially located at the 5’ end of genes overlapping gene promoters ^25^. CpG sites are also found in gene bodies and in other regions that are named CpG shores (2 kb regions flanking CGIs), CpG shelves (>2 kb regions flanking CpG shores), and open sea regions (>4 kb to the nearest CGIs) relative to their proximity to CGIs ^26^. Intergenic regions that are enriched for CpG, but located in open seas, may encompass distal genomic regulatory elements, such as enhancers, silencers and insulators. These transcriptional regulatory elements contain recognition sites for DNA-binding transcription factors, which function either to enhance or repress transcription. Specifically, enhancers are elements able to activate target genes from distal locations independent of their orientation ^27^, while insulators are boundary elements that possess the ability to block or insulate the signals of either enhancers or silencers ^28^. The interplay between protein:DNA complexes defines the spatial organization of the human genome. As a result, chromatin loops are formed, mediated by insulator proteins, such as the CCCTC-binding factor (CTCF), facilitating or blocking enhancer-promoter interactions. However, epigenomic alterations, such as the gain of DNA methylation at the binding site of a regulatory element, could disrupt those interactions ^29^, while removing site-specific DNA methylation has shown to reverse the process thereby providing opportunities for potential clinical therapy ^30^.

Unlike the CpG sites dispersed throughout the genome that are usually methylated, CGIs located at promoter regions are generally unmethylated. In physiological conditions, CGI methylation usually occurs as a mechanism of gene repression in specific regions, such as the inactive X-chromosome, imprinted genes and germline cell-specific genes ^31^. In cancer, DNA methylation becomes aberrant and is mostly characterized by focal hypermethylation around the promoters of genes and gene bodies and global hypomethylation among non-promoter elements ^32,33^ (Figure 1). Promoter hypermethylation is an important mechanism of epigenetic silencing of tumor suppressor genes ^34,35^. DNA methylation in nonpromoter regions is proposed to play a major role in intratumoral expression heterogeneity ^36^. In contrast, global hypomethylation largely affects the intergenic and intronic regions of the genome, which may also result in chromosomal instability. In addition to affecting the gene expression and chromosomal status, aberrant DNA methylation also modulates isoform expression and facilitates mutational events in the adult stem and progenitor cells ^13,33,35^.

**Figure 1.**
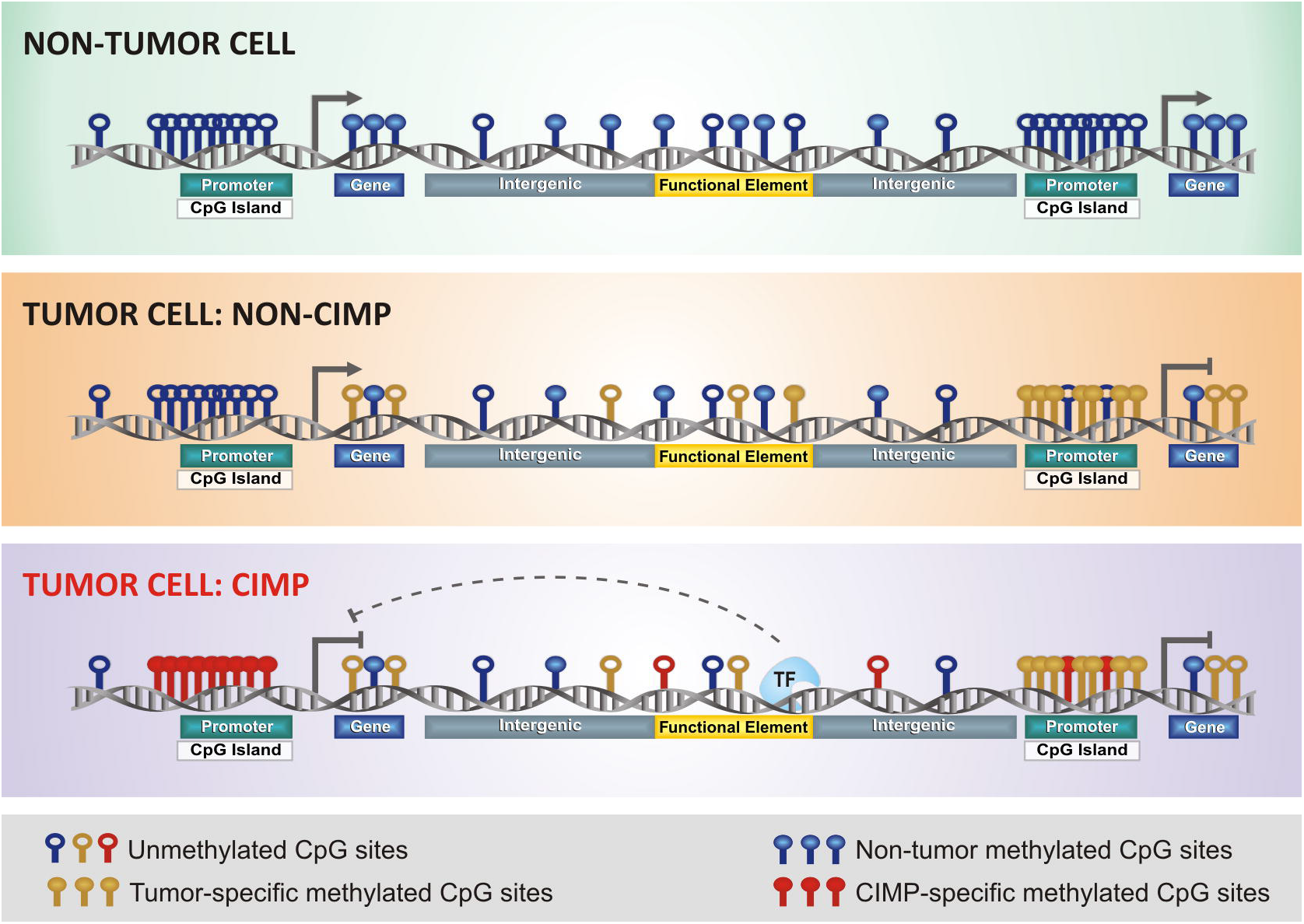
CIMP subtypes in human cancer. This illustration depicts aberrant DNA methylation changes at specific genomic locus in normal and tumor cells, especially in CIMP tumors. Each DNA strand represents one individual methylome. Methylated CpG sites in normal state are represented in blue; non-CIMP tumor DNA methylation gain in yellow, and aberrant DNA hypermethylation in CIMP tumors in red (modified from Weisenberg 2014).

### CIMP: CpG island methylator phenotype

Classically, CIMP is defined by genome-wide hypermethylation of CGI (Figure 1). Tumors carrying this phenotype were first described and validated in the context of colorectal cancers, also known as colorectal CIMP ^37,38^. Since then, CIMP has been described in several tumors, including gliomas ^18,39^. Interestingly, the CIMP+ tumor subset exhibits distinct epidemiological, clinicopathological and molecular features in relation to its CIMP-counterpart. CIMP may have prognostic significance (both favorable and unfavorable) in terms of the OS or predictive value of therapeutic response in certain tumors_^39-41^. Despite tissue-specific differences, evidence suggests that CIMP may be a universal feature across different tumors. Addressing this issue, researchers applied a genome-wide unbiased and unsupervised hierarchical clustering of cancer-specific methylated CGI genes across 15 tumors types; however, gliomas were not included in the cohort ^42^. These researchers observed a set of 89 discriminative loci that allowed the segregation of 12 tumor types into CIMP+ and CIMP-subgroups according to the highest or lowest average levels of DNA methylation, respectively ^42^.

## Glioma Epigenomic Molecular Signatures

### G-CIMP: Glioma-CpG island methylator phenotype

The glioma-CIMP (G-CIMP) subtype was first described by Noushmehr *et al*. in GBM and was further validated in lower-grade glioma (LGG) ^18,19^. The G-CIMP+ subtype occurred frequently in LGG specimens, whereas in GBM it was mostly associated with secondary or recurrent (treated) tumors harboring mutations of the **IDH** gene ^18,19^. By integrating the DNA methylation data with four gene expression clusters previously described in GBM by Verhaak *et al*. (i.e., proneural, neural, classical, and mesenchymal) ^43,44^, the authors found that G-CIMP+ subtypes were highly enriched among the proneural subtypes and in younger patients, compared to G-CIMP negative tumors. Later, the G-CIMP subtype was also described in pediatric glioma patients ^45^. Importantly, G-CIMP+ was shown to be closely associated with *IDH*-mutant gliomas in several studies ^18,19^.

By performing a large multi-platform genomic analysis across 1,122 lower-and high-grade primary adult gliomas, our team, as part of Ceccarelli *et al*., uncovered 7 discrete subtypes with distinct biological features and clinical outcomes ^7^. This DNA methylation-based classification refined and recapitulated previous glioma stratification subgroups based on **IDH** mutation and 1p/19q codeletion status. The subgroups were mainly driven by **IDH*1/2* mutation status and classified as 1: *IDH*-wildtype, enriched for GBM. These were further segregated as classic-like, mesenchymal-like, LGm6-GBM, and PA-like subgroups and 2: *IDH*-mutants, enriched for LGG, and further clustered as 1p/19q codel and non-codel. The *IDH*-mutant non-codel cluster was further refined into two distinct subgroups, based on the extent of DNA methylation, G-CIMP-low (6% of *IDH*-mutant) and G-CIMP-high (55% of *IDH*-mutant), which were determined by a low or high degree of DNA methylation, respectively ^7^.

### Prognostic value of G-CIMP+ and its subsets

The G-CIMP DNA methylation showed relevant prognostic value, independent of the known OS predictors in adult diffuse glioma, such as grade and age ^18^. The favorable prognostic value of G-CIMP+ in both LGGs and GBMs has been reported in many other studies ^7,18,19,45-47^ Notably, *IDH*-mutant G-CIMP+ GBM presented favorable survival and molecular similarities to LGG. On the other hand, LGG carrying *IDH*-wildtype G-CIMP negative subsets presented molecular and clinical behavior similar to GBM ^7,19,45,46^. The refinement of the G-CIMP+ stratification revealed that not all *IDH*-mutant G-CIMP+ tumors had the same prognosis. The G-CIMP-low subset had the poorest OS among the *IDH*-mutant gliomas (median survival G-CIMP-low=2.7 years, G-CIMP-high=7.2 years, cox-regression p<0.001; and vs codel=7.9 years, p<0.001) resembling the behavior observed in the *IDH*-wildtype gliomas (median survival 1.2 years) ^7^.

### Relationship between G-CIMP+/subsets and established prognostic and/or predictive molecular biomarkers

**IDH** mutations, codeletion of 1p/19q, *MGMT* promoter methylation, and G-CIMP+ are all independent favorable prognostic biomarkers ^48^. However, combining some of them has been shown to improve their individual prognostic value ^7,18,48^. For instance, patients harboring a triple combination of 1p19q codeletion, **IDH** mutation and *MGMT* methylation had significantly better OS than those carrying the *MGMT* methylation biomarker alone ^49^. In addition, some of them (codeletion of *1p19q* and *MGMT* promoter methylation) have also been established as predictive biomarkers and have been used in decision making for chemotherapy and/or radiation treatment ^50,51^. The relationship between G-CIMP and the aforementioned biomarkers will be explored in the following sections.

#### **IDH** mutation

One of the most relevant clinical discoveries in neuro-oncology involves the role of **IDH*1* mutations as a prognostic marker and potential as a drug target in glioma ^1,7,10,19,45,52-54^ The R132H **IDH*1* mutation, an amino acid substitution at a single arginine residue in the active site of the enzyme, is highly prevalent in grade II and III gliomas and appears in secondary GBMs, which develop from lower-grade tumors ^10^. Although considerably less common, mutations in **IDH*2*, the mitochondrial homolog of the cytosolic **IDH*1*, have also been identified in gliomas ^55^.

Due to the close relationship between **IDH** mutations and G-CIMP+, it was suggested that the G-CIMP+ prognostic value was mainly due to its relation to **IDH** mutations ^52^. The mechanisms associated with favorable prognosis in *IDH*-mutant G-CIMP+ tumors are still under investigation ^13,18^. However, a meta-analysis of a G-CIMP gene list with prior gene expression analyses suggested that G-CIMP+ tumors may be less aggressive because of the silencing of key mesenchymal genes ^18^.

The predictive value of **IDH** mutations for treatment response is still controversial ^4,50,51,56^; however, a phase III clinical trial reported that *IDH*-mutant anaplastic gliomas benefited from an early combination of procarbazine, lomustine, and vincristine ^51^. The predictive value of G-CIMP+ and its subsets is still unknown.

#### 1p/19q

The 1p/19q codel has an established favorable prognostic and predictive value in gliomas ^47,50^ and is found exclusively in *IDH*-mutant tumors ^57^. The interaction between 1p/19q status and G-CIMP+ has also been reported ^12,58^. The *IDH*-mutant G-CIMP+ codel tumors, for example, were associated with a better OS than the *IDH*-mutant G-CIMP+ non-codel (mean survival G-CIMP+ codel=9.9 years and G-CIMP+ non-codel=4.4 years) ^58^. In our pan-glioma cohort, 9.1% of the *IDH*-mutant G-CIMP+ non-codel subgroup was represented by the G-CIMP-low subtype with the remainder represented by the G-CIMP-high subtype ^7^. As described, among the *IDH*-mutant G-CIMP+ non-codel tumors, the G-CIMP-high subset had a similar survival to the *IDH*-mutant codel tumors ^7^.

#### MGMT promoter methylation

The epigenetic silencing of O-6-methylguanine DNA methyltransferase *(MGMT)* by methylation of its promoter has an established prognostic and predictive value in gliomas and is used in therapeutic decision planning ^14-17^. The MGMT enzyme repairs the DNA damage caused by alkylating agents, such as TMZ, used to treat patients with GBM, playing a key role in tumor cell resistance to the cytotoxic effect of alkylating agents.

*MGMT* promoter methylation is associated with **IDH** mutation and G-CIMP status, regardless of glioma grade ^7,11,59-62^. In fact, the prognostic-versus-predictive value of *MGMT* promoter methylation was found to be at least partly dependent on the context of **IDH** mutation and G-CIMP status. In *IDH*-wildtype gliomas, also considered G-CIMP negative tumors, the *MGMT* promoter methylation was found to be a predictive marker of a favorable response to alkylating agent chemotherapy ^16,50^. In contrast, in gliomas harboring *IDH*-mutation, including G-CIMP+ cases, it was reported that *MGMT* promoter methylation is a prognostic indicator for better survival regardless of treatment with radiotherapy and alkylating agent chemotherapy or with radiotherapy only, but it did not predict the response to treatment ^16,50^.

From our published data^7^, we estimated that among the TCGA *IDH*-mutant cohort, 91.8% of glioma specimens presented with *MGMT* promoter methylation compared to 40.0% among *IDH*-wildtype tumors (p < 0.001). Notably, G-CIMP-low specimens presented a lower proportion of *MGMT* promoter methylation (68.0%) compared to G-CIMP-high specimens (88.8%, p = 0.008, unpublished data). These data suggest that G-CIMP status (-low or-high), as well as **IDH*1/2* mutation, should be considered when determining the predictive significance of *MGMT* promoter methylation in gliomas in revealing the benefit from alkylating chemotherapy.

## Potential Drivers Associated with G-CIMP

Putative driver mechanisms underpinning the aberrant CpG methylation that occurs in CIMP tumors are still under surveillance in gliomas. Several potential cancer-specific mutated driver genes include **IDH*1/2* and *H3F3A* ^18,19,45,63^. Hotspot mutations in **IDH*1/2* genes provoke *IDH* to display a neomorphic enzymatic activity that leads to the reduction of α-KG to 2-hydroxyglutarate (2-HG), an oncometabolite that functions as a competitive inhibitor of α-KG ^54,64,65^. The accumulation of 2-HG impairs the activity of α-KG-dependent-dioxygenases, such as histone and DNA demethylases (e.g., TET enzymes), leading to global hypermethylation (CIMP), as well as the impairment of cell differentiation ^19,64,66^. Another consequence of **IDH** mutations is the alteration of the intracellular levels of the coenzyme NAD+ ^67^. NAD+ has a key role in intracellular signaling pathways implicated in cancer cell growth. It was shown that *IDH*-mutant cells had a reduced expression of NAPRT1 (nicotinate phosphoribosyl transferase 1), a rate-limiting enzyme within the NAD+ salvage system. Reduced expression of NAPRT1 led to lower basal NAD+ levels, rendering the *IDH*-mutant cell more vulnerable to death ^67,68^. Furthermore, **IDH** mutations also result in the methylation of histones that contribute to DNA methylation by themselves ^69^. Histone *H3F3A* mutations were also reported to drive the global aberrant pattern of DNA methylation in a subset of pediatric GBM by as yet unknown mechanisms ^45^. Interestingly, in a subset of GBM, the authors found recurrent and mutually exclusive mutations either in **IDH*1* or *H3F3A* (affecting amino acids K27 or G34) with distinct clinical, genomic, and epigenomic features ^45^.

Recent large-scale genome-wide association studies (GWAS) provided evidence for a defined germline variant located on chromosome 9p21.3 which was found to be enriched among G-CIMP tumors ^70^ and a variant mapped to an intronic region located on chromosome 2q33.3, 50K base pairs from **IDH*1* ^71^. Despite the lack of functional experiments, these findings, in combination with known somatic alterations, offer potential insights on the role of genetic variants in the biology and etiology of G-CIMP tumor development and possibly progression.

Figure 2 summarizes the major milestones in integrating genomics and epigenomics data to uncover glioma molecular and clinical phenotypes that led to the characterization (either directly or indirectly) of G-CIMP subsets.

**Figure 2.**
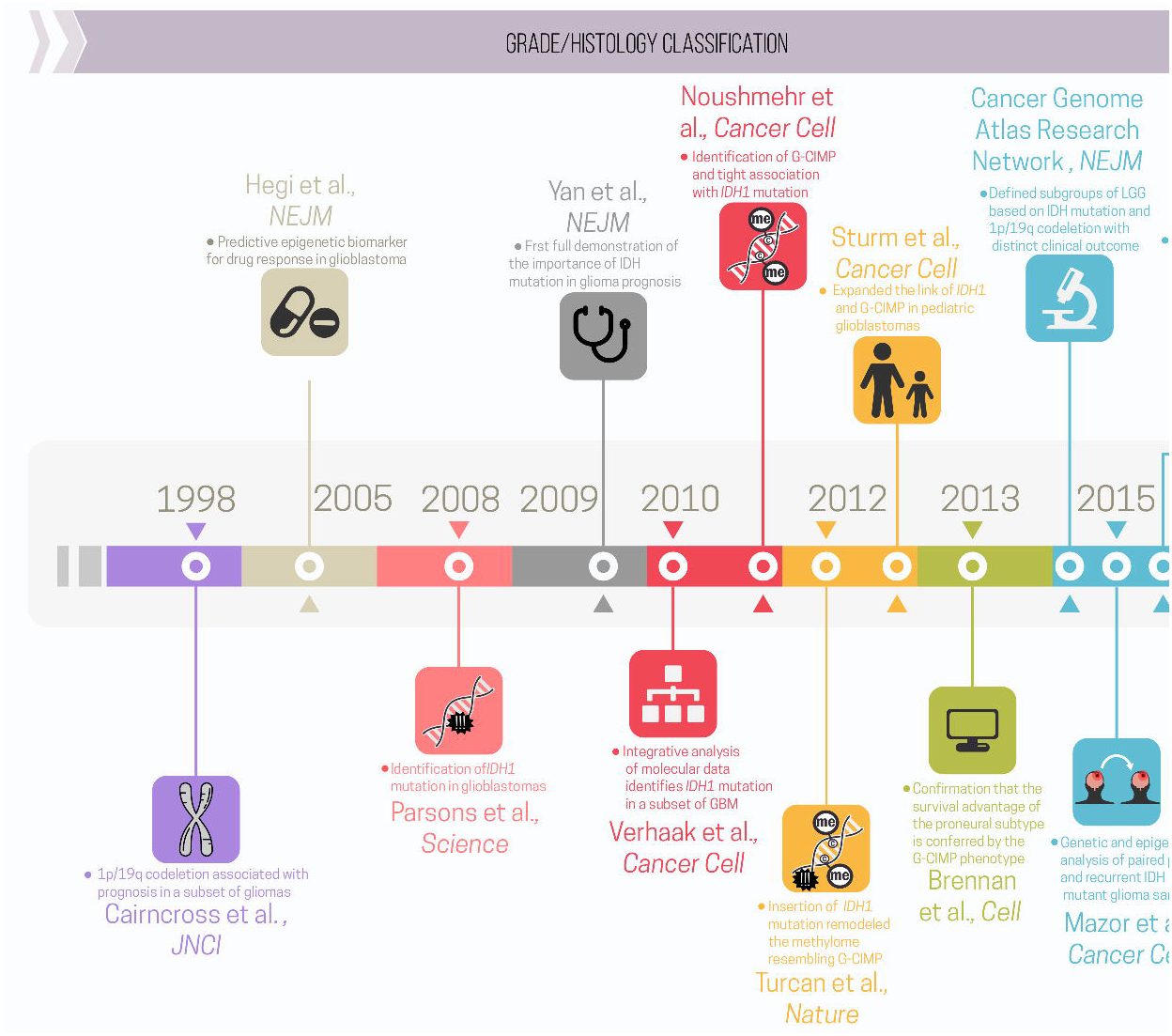
Summary of major milestones in the integration of large-scale genomic and epigenomic data that uncovered molecular and clinical features associated with pediatric and adult gliomas (timeline). Each milestone is indicated by marker papers that reported key molecular findings with clinical implications, along with a bullet summarizing their contribution. The timeline is broken up by the 2016 WHO publication (before and after) (Modified from the original copy-free design by Freepik).

## G-CIMP+ Subsets and Glioma Progression/Recurrence

WHO grades II and III **IDH**-mutant non-codel (astrocytic) gliomas habitually recur and unpredictably undergo malignant transformation to highly aggressive and treatment-resistant grade IV GBM ^1,72^. Remarkably, the analysis of a small cohort of primary and recurrent matched tumor specimens, composed of LGG and GBM, revealed that some G-CIMP-high tumors exhibited a demethylated pattern after relapse that was similar to those observed in the G-CIMP-low gliomas, suggesting a progression from the G-CIMP-high to the G-CIMP-low subset ^7^. Unpublished work from our own laboratory may provide further refinement of the epigenetic shift from G-CIMP-high to G-CIMP-low upon tumor recurrence in TCGA and non-TCGA specimens ^73^. Interestingly, in that cohort, G-CIMP-low at recurrence appeared in 12% of all gliomas and shared epigenomic features resembling *IDH*-wildtype primary GBM ^73^. Genome-wide decreases in DNA methylation levels associated with progression have also been reported by other studies ^74,75^.

The hypothesis that G-CIMP-high tumors may relapse as G-CIMP-low gliomas suggests that variations in DNA methylation could be a key determinant of the mechanisms that drive glioma progression. Notably, the majority of CpG sites that underwent significant DNA demethylation in G-CIMP-low recurrent tumors were primarily found within intergenic (open sea) regions ^7^ (Figure 1). CTCF, a methylation-sensitive insulator protein, has an important role in stabilizing enhancer-gene interactions in intergenic regions, as mentioned previously in this review. In *IDH*-mutant gliomas, it was shown that hypermethylation at CTCF binding sites reduced CTCF binding. The consequent loss of insulation led to aberrant enhancer-gene interactions that ultimately resulted in the upregulation of a glioma oncogene ^76^. Since the discovery of *IDH*-mutant glioma subtypes, our group began to investigate the potential role of DNA methylation status at a single-base pair resolution using Whole-Genome Bisulfite Sequencing (WGBS). Confirming previous data ^7^, our unpublished findings revealed that, compared to G-CIMP-high (as well as to non-tumoral brain specimens), G-CIMP-low presented DNA hypomethylation at some CTCF binding sites. Collectively, these findings may offer potential support for the hypothesis that the loss of DNA methylation at CTCF binding sites will also influence the chromatin architecture, which is mediated by the disruption of insulator binding. This phenomenon will, in turn, dysregulate nearby genes (Figure 3) (Sabedot TS et al, unpublished data). However, further studies are needed to confirm this hypothesis. We also found that the hypomethylated intergenic regions were enriched for the OLIG2 and SOX-family binding motifs ^7^, which have been described as neurodevelopmental transcription factors essential for GBM propagation ^77^. SOX-family genes are transcription factors that are also involved in the induction and maintenance of stem cell pluripotency ^78^, promote self-renewal of neural stem cells in the nervous system ^79^, and regulate the plasticity between glioma stem cell and non-stem cell states within brain tumors ^80^. Accordingly, G-CIMP-low tumors displayed abnormalities in cell cycle pathway genes, such as *CDK4* and *CDKN2A*, supporting the association between stem cell signaling pathways and tumor proliferation in gliomas. Therefore, loss of CpG methylation at these functional genomic elements, known to be associated with normal development and pluripotency, defines a possible mechanism of glioma progression ^7,73^.

**Figure 3.**
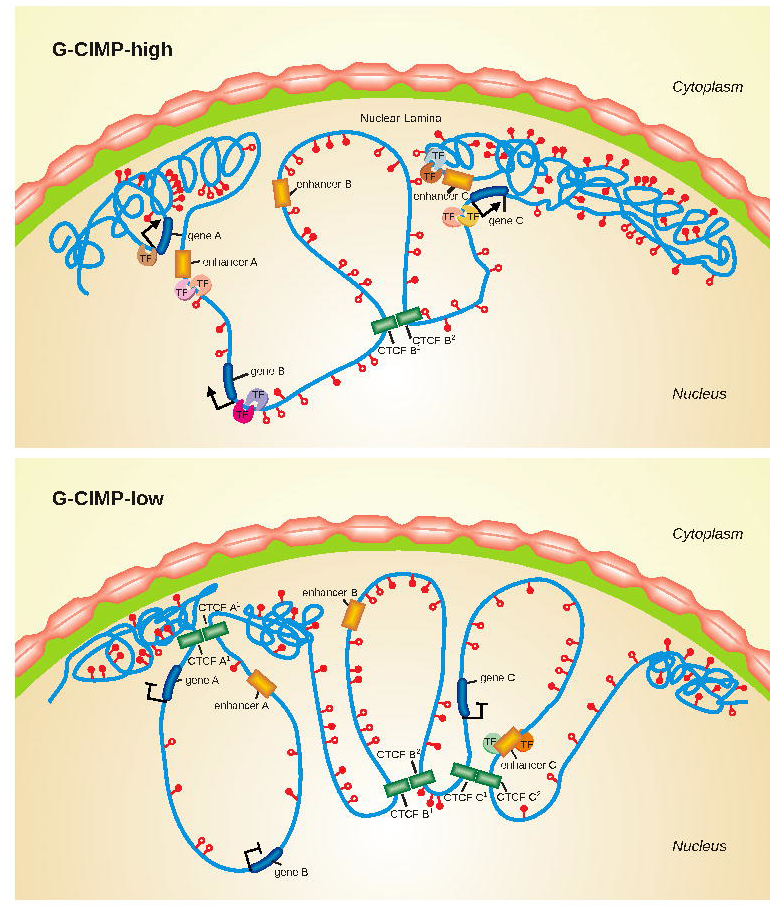
Chromatin changes in the progression of G-CIMP-high to G-CIMP-low tumors. This illustration shows a model of chromatin reorganization during the progression from G-CIMP-high to G-CIMP-low. G-CIMP-low (lower panel) shows a loss of DNA methylation at specific loci causing disruption of CTCF binding sites, reorganization of chromatin, and dysregulation of gene expression (upper panel).

In addition, Bai *et al*. found that key developmental transcription factors that are regulated by Polycomb Repressive Complex 2 (PRC2) in human stem cells became hypermethylated during glioma progression resulting in gene silencing, which could ultimately cause GBM cells to enter a continuous state of stem cell-like self-renewal ^75^. Mazor *et al*. reported specific hypomethylation events that may contribute to the increased cell proliferation upon progression from LGGs to GBMs ^74^. Recently, we derived a metric to measure the degree of de-differentiation of tumors based on DNA methylation and found a strong association between a high undifferentiated score and glioma molecular subtypes associated with worse clinical outcomes (i.e., G-CIMP-low, classic-like, mesenchymal-like, LGm6-GBM, and PA-like). Interestingly, G-CIMP-low patients showed higher undifferentiated scores compared to G-CIMP-high patients, resembling those found in primary *IDH*-wildtype tumors (Malta TM et al, unpublished). This finding may suggest that a stem cell-like phenotype may be involved in unfavorable clinical outcomes and reinforce the importance of epigenetic alterations occurring in intergenic regions that may disrupt important regulatory elements affecting oncogenesis and tumor progression (Figure 4).

**Figure 4.**
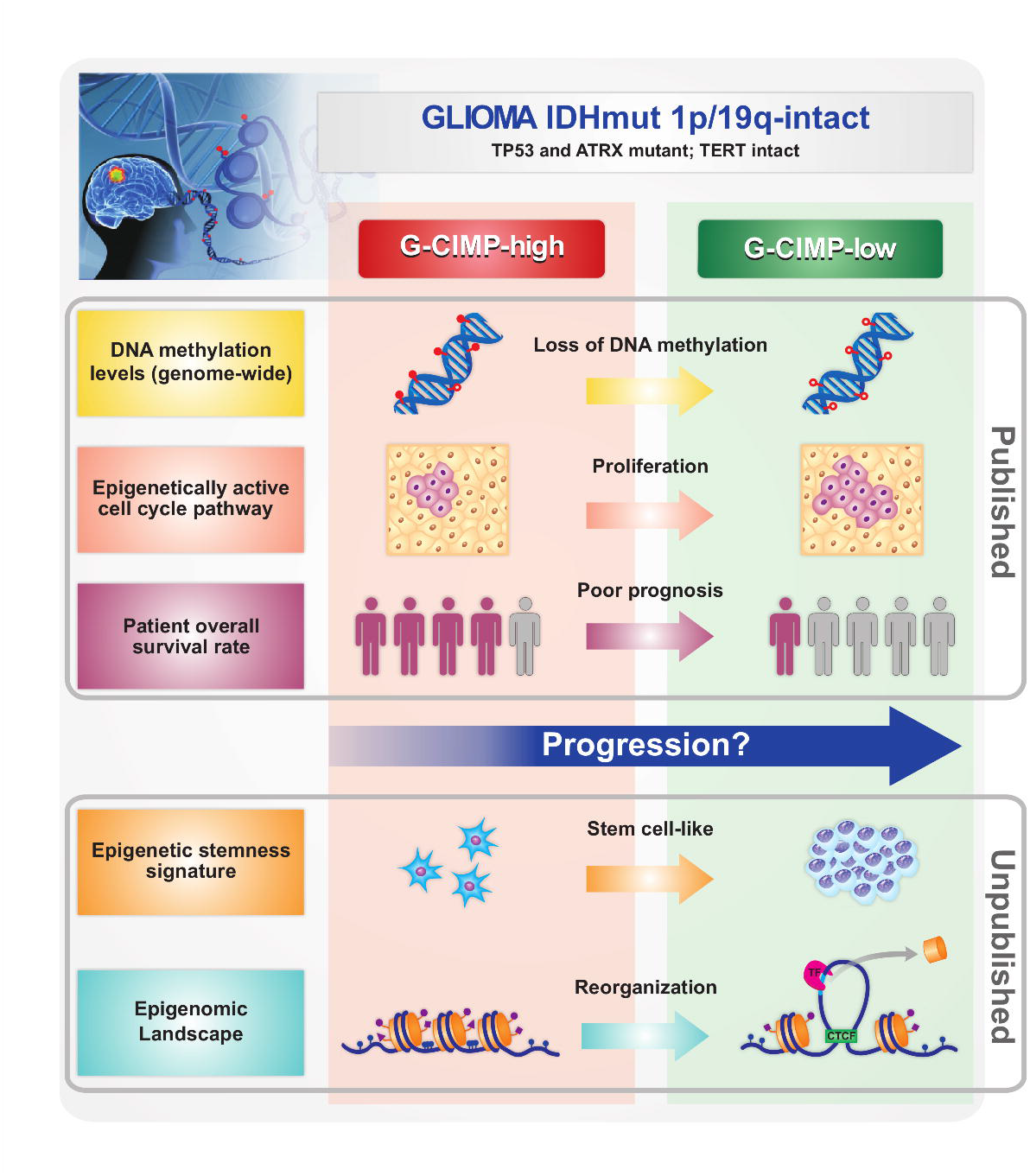
Overview of major discoveries that define G-CIMP-high and G-CIMP-low glioma subsets. G-CIMP-high and G-CIMP-low tumors share the following genomic alterations: *IDH* mutant-1p/19q intact, TERT promoter wild type, and ATRX and TP53 mutant. However, the G-CIMP-low subset defined a subgroup of *IDH*-mutants 1p/19q intact gliomas associated with DNA demethylation. Changes in chromatin architecture led to an imbalance between the insulators and enhancers and the consequent activation of cell-cycle related genes, the increase in stemness features, and poor clinical outcome compared to G-CIMP-high gliomas (cartoon representation, not to scale).

## G-CIMP Detection

The diagnosis of G-CIMP was reported by Noushmehr *et al*., 2010 ^18^. This diagnosis was based on an epigenetic biomarker panel consisting of seven hypermethylated loci *(ANKRD43, HFE, MAL, LGALS3, FAS-1, FAS-2*, and *RHO-F)* and one hypomethylated locus, *DOCK5*, validated *in silico*. A sample was considered G-CIMP+ if at least six genes displayed a combination of *DOCK5* DNA hypomethylation and/or hypermethylation of the remaining genes in the panel. The utilization of the MethyLight assay to detect the aforementioned biomarkers showed perfect concordance with the results obtained by array platforms and may be suitable for clinical utilization ^18^.

Later, G-CIMP-high and-low subsets were defined and validated *in silico* by a panel of 163 DNA-methylated probe signatures using methylation arrays ^7^. Our group developed a concise DNA methylation biomarker panel, derived from those probes, consisting of 14 methylated probes allowing the identification of the seven glioma subgroups that were previously reported ^7^ and validated *in silico* (unpublished data). Eight of those probes distinguished between G-CIMP+ subsets (-low and-high) in the context of *IDH*-mutant glioma specimens; however, their detection by more readily available assays warrants investigation.

## Epigenomic-and Chromatin-Targeted Therapies

Epigenetic mechanisms are heritable and potentially actionable, making them an attractive target for the treatment of diseases, including cancer ^81^. Many of the epigenome-targeting drugs have been demonstrated to be beneficial and safe in hematological malignancies ^82,83^. However, studies in solid tumors are limited ^53,81,84^. Epigenetic drugs approved or in clinical trials have been thoroughly explored in recently published reviews ^53,81,85^.

In gliomas, few trials from this class have been completed ^82,83,85,86^. To our knowledge, there are no clinical trials specifically addressing G-CIMP+ subtypes. However, preclinical and clinical studies using drugs targeting DNA methylation directly, or the mechanisms involved in G-CIMP, are underway. Hypomethylating agents such as the inhibitors of DNA methyltransferase (DNMT), the histone deacetylase (HDAC) and the bromodomain and extra-terminal motif proteins (e.g., BRD4 inhibitor) represent epigenetic drugs with broad actions ^81^.

DNMT inhibitors or DNA demethylating agents (e.g., 5-Azacytidine, 5-Aza-2-deoxycytidine-decitabine) promote dose-dependent global demethylation activity by depleting or degrading DNA methyltransferases. This may reverse aberrant expression of genes related to nearly all pathways involved in cancer initiation and progression (i.e., tumor suppressor genes, oncogenes, genes associated with stemness or pluripotency, apoptosis, cell-cycle and, immune response) ^87,88^. Moreover, DNMT inhibitors were also shown to reverse tumor immune evasion through viral infection mimicry and up-regulation of viral defense pathways ^89-91^. For instance, in an **IDH*1*-mutant glioma xenograft, low doses of the demethylating agent, decitabine, restored the activity of DNMT1 and, consequently, reversed DNA methylation marks in promoters of differentiating genes. This change resulted in the loss of stem-like features and in the arrest of glioma growth in that model ^92^. Although preclinical studies using these drugs in gliomas seem promising, a corresponding clinical trial using decitabine on adult glioma has not been reported. Considering the potential adverse impact of loss of methylation in the progression or recurrence of gliomas, reported previously ^73,74^, it seems reasonable to take G-CIMP subsets into account during clinical trial design.

Targeting **IDH**-mutant tumors, including G-CIMP+ gliomas, with the **IDH**-mutant enzyme inhibiting agents is currently under investigation in clinical trials ^53,67,81^. The mechanistic intent of these drugs involves the metabolic pathways altered in *IDH*-mutant tumors (i.e., decrease of 2-HG in hematological or salvage of NAD+ in solid tumors) ^61^. However, a preclinical study has not identified a decrease or epigenetic change in *IDH*-mutant cancer initiating cells exposed to an *IDH*-R132 inhibitor drug, despite marked reduction in 2-HG ^67^. Tateishi et al. postulated that the activity of this class of drugs may be limited to a subset of tumors. Moreover, NAD+ metabolic depletion was recently shown to be an attractive therapeutic target in **IDH*1*-mutant cells vulnerable to a cytotoxic response when exposed to a nicotinamide phosphoribosyl transferase (NAMPT) inhibitor. The same response was not seen in **IDH*1-* wildtype cell lines ^67^; however, a later preclinical model demonstrated that NAMPT inhibitors enhanced the cytotoxic effects of TMZ in GBM cells ^93^.

Immune modulation is also affected by epigenomic alterations and has been shown to be an attractive target for pharmacotherapy in cancer ^94^. The identification and targeting of disease-specific neoantigens has demonstrated success in other diseases, renewing excitement for immunotherapy in glioma. An immunogenic epitope vaccine targeting mutant **IDH*1* in glioma with promising preclinical work is now in phase I clinical trials ^95,96^. Several other clinical trials are underway in glioma, including phase III targeted immunotherapy trials ^97^. The use of immune checkpoint inhibitors targeting programmed cell death protein 1 (PD1) and/or its ligand 1 (PD-L1) has also shown activity in several cancer types and is currently being tested in GBM clinical trials. *PD-L1* promoter methylation and lower *PD-L1* expression in *IDH*-mutant gliomas compared to *IDH*-wildtype point to an epigenetic regulation of *PD-L1* and indicate that patients harboring *IDH*-mutant gliomas may not benefit from monotherapy with drugs targeting the blocking of the PD1/PD-L1 pathway ^98^.

Co-administration of epigenomic agents (e.g., DNA-demethylating agents and HDAC inhibitors) has demonstrated improved efficacy of immunotherapy in many tumor types by increasing the tumoral immune-response, enhancing the expression of immune blockade checkpoints, and by reducing cellular adaptation that leads to drug resistance ^99^. Given the potential epigenetic regulation of PD-L1 expression, patients harboring the **IDH**-mutant (G-CIMP+) tumors may benefit from combined treatment modalities, including demethylating agents and PD1/PD-L1 inhibitors. The relationship between the G-CIMP subsets and response to immunotherapy, either as a predictive marker or epigenetic therapy target, is largely unknown and worthy of further investigation.

## Final Remarks and Perspectives

Advances in glioma research have highlighted the significance of epigenomic alterations. The incorporation of genetic markers into the traditional WHO histopathological classification of CNS tumors reflects widespread adoption of the latest scientific and clinical advances in molecular neuro-oncology into clinical practice ^1^. Moreover, evidence suggests that comprehensive analysis, such as unsupervised glioma subtyping based on gene expression or on G-CIMP status, rather than individual molecular markers, may improve prognostic and predictive outcomes ^100^. Currently, the guidelines for the detection of established prognostic biomarkers are based on individual marker assays: for example, immunohistochemistry or DNA sequencing for mutations in **IDH*1/2* and *H3F3A* genes, fluorescent in situ hybridization or microsatellite PCR-based loss of heterozygosity analyses for codeletion of chromosomal arms 1p and 19q, and real-time methylation-specific PCR for *MGMT* promoter methylation ^61^. However, the advent of genome-wide analysis technologies has enabled the concurrent detection of DNA methylation patterns and genomic and copy number alterations in tumor specimens ^61^. Remarkably, the identification of a DNA methylation-based classifier of glioma specimens enabled researchers to capture already known prognostic markers, such as **IDH** mutation and 1p19q codeletion ^7^. In addition, it uniquely enabled the identification of a subset of tumors within the *IDH*-mutant/G-CIMP+ group (i.e., G-CIMP-low) that presented a prognostic disadvantage in relation to their counterparts (*IDH*-mutant G-CIMP-high non-codels and codels). In addition to refining glioma classification, the identification of G-CIMP+ subsets also shed light on the role of the demethylation of specific genes in the pathogenesis of glioma progression, independently of grade or **IDH** status ^7^.

In summary, the detection of G-CIMP+ and its subsets (-high and-low) builds on the 2016 WHO molecular effort to refine glioma classification (Figure 5) and provides insight into disease course and treatment opportunity ^73^. The prognostic significance of G-CIMP+ across all glioma types has been confirmed in many studies ^7,18,19,45-47^ It is possible that, as for other established therapeutic predictive markers in gliomas, such as 1p/19q codeletion and *MGMT* promoter methylation, G-CIMP+ subsets will be used for patient counseling and be part of algorithms used for clinical trial design and for therapeutic decisions ^4^. However, the utility of these classifiers and biomarkers in planning treatment strategies and designing clinical trials has not been fully validated to date.

**Figure 5.**
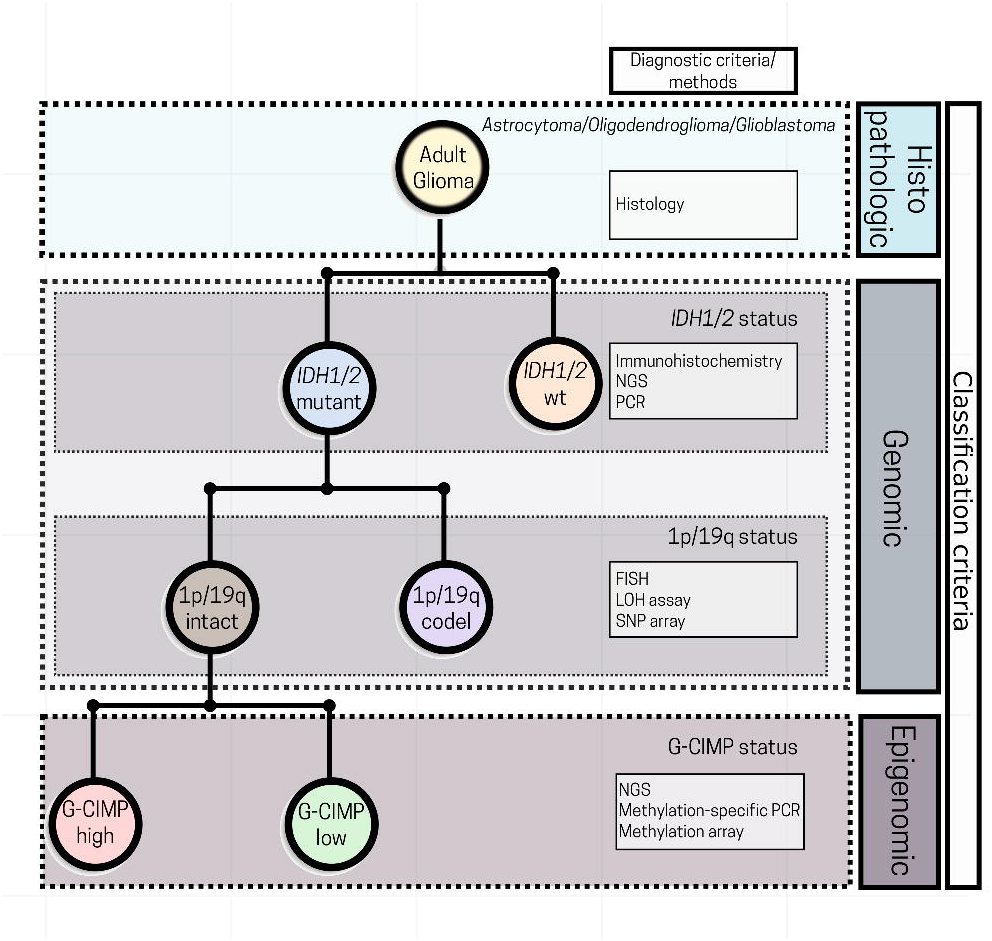
Perspectives in incorporating G-CIMP subsets into the current 2016 WHO glioma classification algorithm. A simplified diagram for glioma classification based on histological, genetic and epigenetic features. The incorporation of G-CIMP subsets further refined glioma classification. NGS: next-generation sequencing; PCR: polymerase chain reaction; FISH: fluorescent in situ hybridization; LOH: loss of heterozygosity; SNP: single-nucleotide polymorphism; wt: wildtype (Modified from the original copy-free design by Freepik).

## Acknowledgments

The authors thank Sandra Navarro at the Regional Blood Bank of Ribeirao Preto for carefully assembling the figures and OMICs laboratory members for their helpful discussion and contribution. We thank Michelle Felicella and Chunhai (Charlie) Hao for insightful discussion about glioma classification. We would also like to thank Susan MacPhee-Gray, Ana deCarvalho, Laila Poisson and Tobias Walbert for critical review of the manuscript.

